# Uptake mechanisms and physiological effects of furanic compounds from the Maillard reaction in budding yeast

**DOI:** 10.1101/2025.08.08.669114

**Authors:** Laise Cedraz Pinto Matos, Amy Milburn, Chris MacDonald

## Abstract

Maillard reaction products (MRPs) are formed during the thermal processing of foods and exhibit important sensory attributes. Furanic compounds are a subset of MRPs commonly found in food products that are toxic to eukarytoic cells, although the mechanisms of toxicity are poorly understood. We used budding yeast to explore uptake mechanisms of common furanic compounds: 5-hydroxymethylfurfural (HMF), furfural (FUR), and 2-Furyl methyl ketone (FMK). Titrations of each furanic compound were used to identify concentrations that have an inhibitory effect on growth. We identified HMF as a potential substrate of the Pdr5 multidrug resistance pump and linked HMF and FUR toxicity to surface nutrient transporter levels. Live cell imaging shows that HMF disrupts mitochondria whilst FUR affects the endolysosomal system. Results indicate these furanic compounds may have distinct uptake, efflux, and toxicity mechanisms. As many of these cellular components are conserved throughout evolution, this work could shed light on the metabolism of toxic compounds commonly found within animal food sources.

## INTRODUCTION

The Maillard reaction is a complex cascade of non-enzymatic browning processes between reducing sugars and amino acids, and is fundamental to the development of flavour, aroma, and colour in thermally processed foods (Liu et al., 2022). The array of molecules produced, collectively termed as Maillard reaction products (MRPs), have distinct sensory implications for the food industry (Hosry et al., 2025; Tamanna & Mahmood, 2015). The interplay of various factors, such as chemical composition, temperature, time, humidity, and pH, along with the presence of glycation agents or oxidants, dictates the progression of this chemical cascade and the specific MRPs generated (Nie et al., 2013; Shakoor et al., 2022). Various MRPs have been identified in foods, common examples include acrylamide and furanic compounds (Agcam, 2022; Sun et al., 2022). Humans are frequently exposed to furanic compounds, such as 5-hydroxymethylfurfural (HMF), furfural (FUR), and 2-Furyl methyl ketone (FMK), which are associated with sugar-rich foods (National Toxicology Program, 2010). However, beyond sensory appeal, this reaction also generates a broad spectrum of potentially harmful by-products (Kathuria et al., 2023).

The toxicity of MRPs has been extensively debated, with examples like acrylamide being correlated with various diseases and negative effects (Başaran et al., 2023). However, biological effects of MRPs are context-dependent, varying according to factors like dose, stability, biological model, and metabolic capacity, (Cheng et al., 2022; Wierckx et al., 2011). Furanic compounds are known to be carcinogenic (National Toxicology Program, 1993). Studies using rodent species have shown that ingestion of furanic compounds induces DNA damage on spleen/liver cells, triggering chromosomal aberrations and cell death (Leopardi et al., 2010; Neuwirth et al., 2012; Yilmaz et al., 2023). In addition to genotoxic effects, furanic compounds are known to induce oxidative stress and inflammation (EFSA Panel et al., 2017). For example, HMF increases reactive oxygen species (ROS) in both fly larvae and cultured human cell models, with the latter example showing ROS triggers apoptosis via mitochondrial pathways (Chen et al., 2024; Qiu et al., 2022). Furanic compounds are known to be produced in substrates used for microorganism conversion to biofuel, which hinders the fermentation process (Wang et al., 2016). In *Saccharomyces cerevisiae*, exposure to these compounds is toxic and leads to elevated ROS levels (Allen et al., 2010). This oxidative stress response may be linked to mitochondrial dysfunction, as directed-evolution approaches aimed at increasing furanic tolerance consistently recovered mutations affecting mitochondrial pathways (Ren et al., 2024).

MRPs are present in food in free form or conjugated to proteins, peptides or polyphenols (Nowak et al., 2021; Zhu et al., 2020). The health effects of these compounds are primarily evaluated in isolation, as MRPs are readily absorbed in the gastrointestinal tract and thus these forms represent the bioavailable fraction (Pagare et al., 2024). Regulatory agencies use measurements of compounds such as HMF, acrylamide and furfurals in free form to estimate dietary exposure, construct safety limits and compare with the acceptable daily intake (EFSA_Panel et al., 2017; Eisenbrand, 2020). The ready conversion of bound to free MRP compounds after acid and enzymatic digestion has already been demonstrated in cells (Zeng et al., 2020). Furthermore, the use of isolated analytical standards also allows the evaluation of toxicity and cellular mechanisms with standardized and reproducible protocols (Qiu et al., 2022), meeting the requirements of toxicological evaluation. Thus, the free form of MRPs is more directly related to acute systemic effects, such as cytotoxicity, genotoxicity, and metabolic alterations, which are biologically more relevant.

Despite extensive toxicological characterisation of furanic aldehydes, a longstanding consensus in the literature is that small furan derivatives, such as HMF, FUR, and related molecules, enter eukaryotic cells predominantly by passive diffusion across the lipid bilayer. This view is grounded in their low molecular weight, moderate polarity, and aromaticity, all of which favour membrane permeability without requiring carrier proteins (Heer & Sauer, 2008; Klinke et al., 2004). Consequently, passive diffusion has been widely accepted as the default explanation for cellular uptake of these compounds, even though it does not exclude the possibility that transporter systems modulate their intracellular retention or contribute to compound-specific toxicity. Other studies have challenged the general assumption that xenobiotics freely cross membranes, arguing instead that many small molecules previously considered to be passively permeable are in fact subject to transporter-mediated flux in eukaryotic cells (Baril et al., 2023; Kell, 2021). Revisiting this debate is therefore essential for understanding how furanic inhibitors interact with conserved eukaryotic pathways of stress response, detoxification, and organellar homeostasis.

Eukaryotic cells adjust their uptake and export of nutrients in response to environmental conditions. *Saccharomyces cerevisiae* is a powerful platform for dissecting these mechanisms at the molecular level. Yeast employs a broad array of surface transporters for amino acids, sugars, and metal ions (Bianchi et al., 2019; Donzella et al., 2023). Yeast surface proteins are routinely internalized to endosomes and follow conserved recycling pathways back to the plasma membrane (Laidlaw et al., 2022; MacDonald & Piper, 2016, 2017). Alternatively, internalized surface proteins can be ubiquitinated and sent through the Endosomal Sorting Complexes Required for Transport (ESCRT) mediated pathway to the yeast lysosome (or vacuole) for degradation (Laidlaw & MacDonald, 2018). This downregulation can be triggered in response to substrate or stress, and is mediated upstream of ESCRTs by the arrestin related trafficking adaptors, which specifically unite transporters with the E3 ubiquitin ligase Rsp5 to mediate ubiquitination (C. H. Lin et al., 2008; MacDonald et al., 2020; Nikko et al., 2008). Beyond these traditional nutrient uptake mechanisms, yeast also express multidrug resistance transporter proteins, that transport various substances and toxic compounds, out of the cell (Piecuch & Obłąk, 2014). Amongst these, the Pdr5 multidrug transporter is the best characterized, which has remarkable promiscuity for diverse substrates, and is therefore of interest in clinical and biotechnological applications (Golin & Ambudkar, 2015; Rogers et al., 2001). Whether furanic compounds are substrates of any of these transporters, in yeast or other eukaryotes, has not been directly tested and is not well understood. However, recent transcriptome analyses did suggest expression of pumps like Pdr5 might correlate with toxicity of furanic compounds added to anaerobic yeast cultures of a xylose-utilizing yeast strain (Ask et al., 2013)

Functional aspects of surface transporters are conserved between yeast and mammalian cells (Belle & André, 2001). Additionally, transporter regulatory mechanisms, such as ubiquitin mediated degradation, engagement with alpha arrestins, and endolysosomal trafficking are also evolutionarily conserved (Alvarez, 2008; Laidlaw & MacDonald, 2018; Piper & Katzmann, 2007). In this study, we hypothesise furanic compounds are actively transported across the plasma membrane, and that uptake and downstream cellular effects could be evolutionarily conserved. Using yeast, we use different genetic systems to explore transporter regulation of furanic compounds, and cell biological approaches to assess disruption of cellular processes. This work sets out to define fundamental mechanisms contributing to furanic-induced toxicity and the physiological responses of eukaryotic cells, ultimately contributing to a broader understanding of food-derived toxicants and their impact on human health.

## RESULTS

### MRPs induce toxic effects in yeast cells

To determine if the budding yeast *Saccharomyces cerevisiae* responded to different furanic compounds, we selected three representatives commonly found in food: HMF, FUR and FMK (**Figure 1**). For initial toxicity experiments, we optimised a 96-well plate end-point growth assay, where cell density at the start of the experiment was compared to later time points. Using this assay, we tested titrations of HMF, FUR and FMK across ranges predicted to exert a physiological effect, based on previous studies in mammalian cells (Kong et al., 2019; Qiu et al., 2022; Zhao et al., 2017). These experiments showed that yeast exposed to HMF at concentrations of 125 µg/ml or less had no significant defect in growth after 24 hours (**Figure 2A**). However, at higher HMF concentrations, growth was inhibited in a concentration dependent manner. To validate these results, we employed our recently optimised continuous measurement growth assay and analysis pipeline (Laidlaw et al., 2025), which also showed a HMF concentration dependent inhibition of yeast growth, with reduced growth efficiency observed for 250 µg/ml and above, with almost complete growth inhibition at concentrations of 2000 µg/ml or higher (**Figure 2B**). We performed similar analysis with other MRPs. Both assays revealed a concentration dependent effect on yeast growth in media containing FUR, with significant growth inhibition at concentrations of 125 μg/mL and above (**Figures 2C - 2D**). Similarly, titrations of FMK showed a range of growth effects, with inhibition in the range 125-250 μg/mL and more potent effects at concentrations of 500 μg/mL and above (**Figures 2E - 2F**).

**Figure 1:**
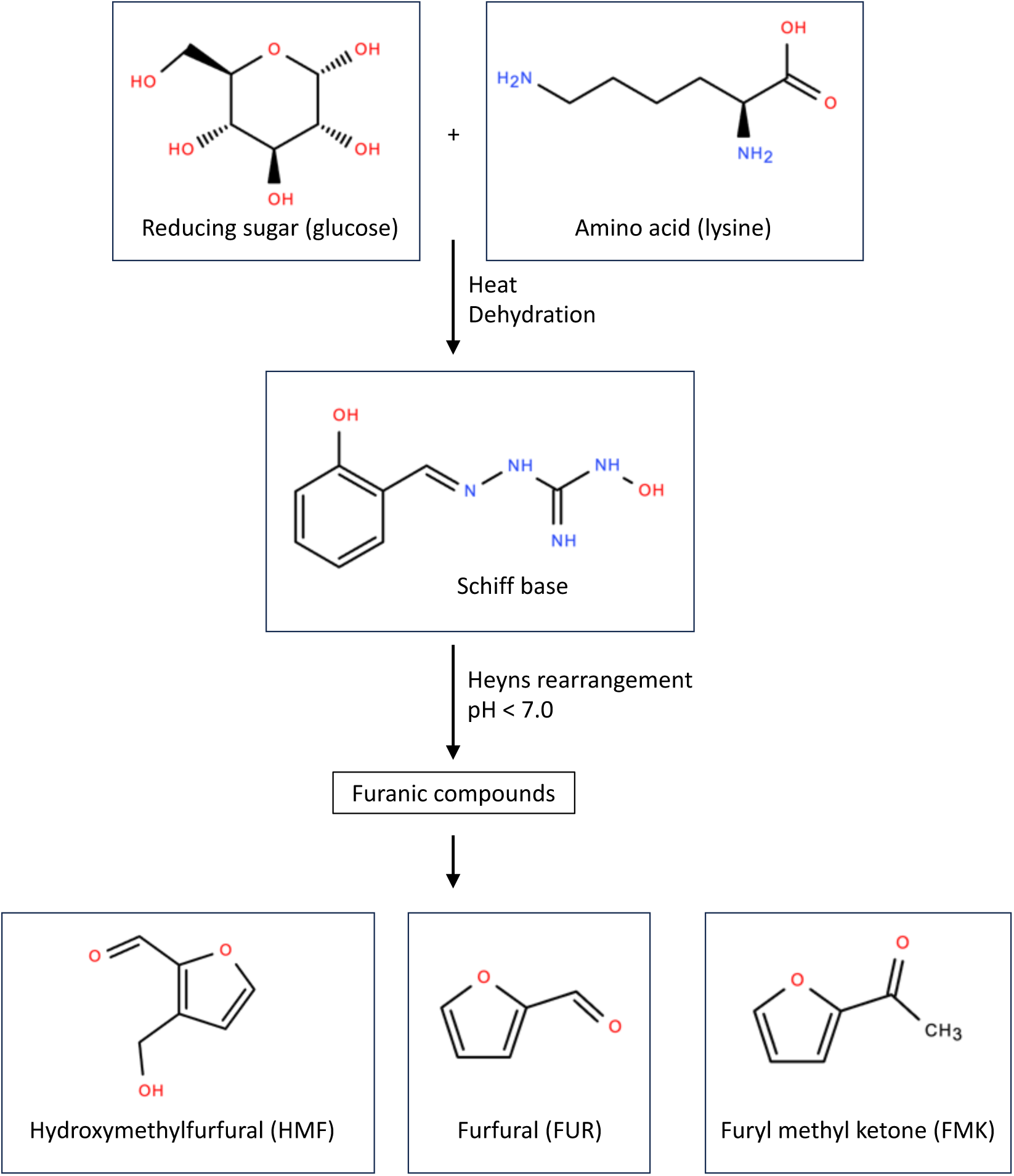
Chemical cascade of the Maillard reaction The non-enzymatic reaction between carbonyl groups of reducing sugars and an amine group triggers a cascade with many products. Those specific to the production of furfurals are depicted.

**Figure 2:**
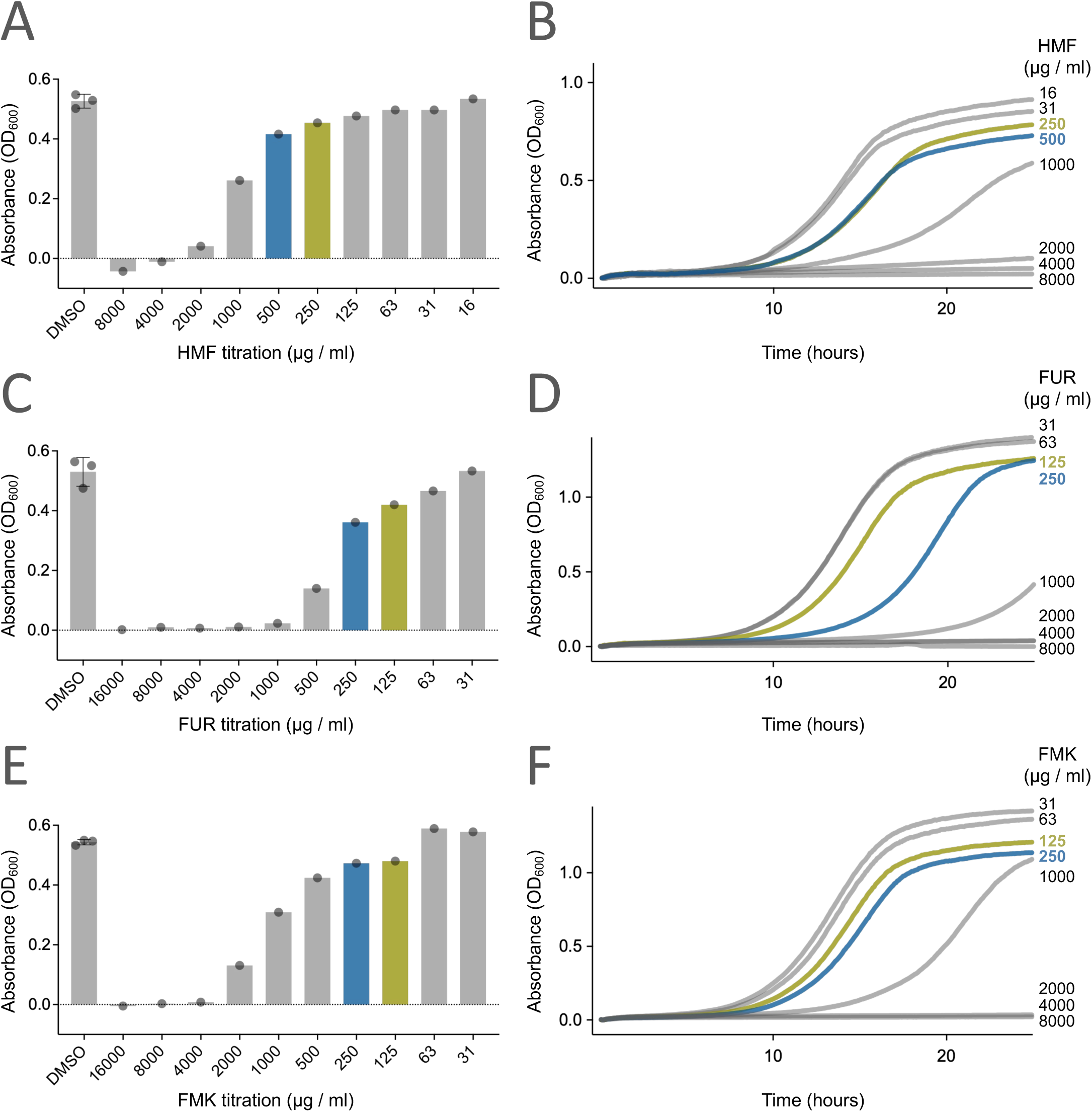
Furanic compounds exert toxic effects on yeast cells **A, C, E)** Endpoint assays were performed with wild-type cells were exposed to varying concentrations of Hydroxymethylfurfural (HMF), Furfural (FUR) and 2-Furyl methyl ketone (FMK). The OD_600_ values were measured of 200 µl cultures grown in 96-well plates immediately after inoculation and then again following 24 hours growth at 30°C. **B, D, F)** Yeast growth assays across a titration of HMF, FUR and FMK were performed in liquid cultures with continuous measurements of OD_600_ every 5 minutes. In each experiment, concentrations of furanic compounds which are used for downstream experiments are indicated (blue and green).

Collectively, this work shows that all three compounds from the Maillard reaction have toxic effects on yeast growth. We therefore set out to use the yeast system to test the hypothesis that transporters influence the cellular uptake and resulting intracellular accumulation of furanic compounds, which are related to common transporter substrates, like sugars and amino acids (Ask et al., 2013; Cong et al., 2021). We reasoned any such non-canonical flux via transporters would not be specific, so sought genetic approaches broadly affecting pathways, instead of specific transporters, to test this hypothesis (**Figure 3**).

**Figure 3:**
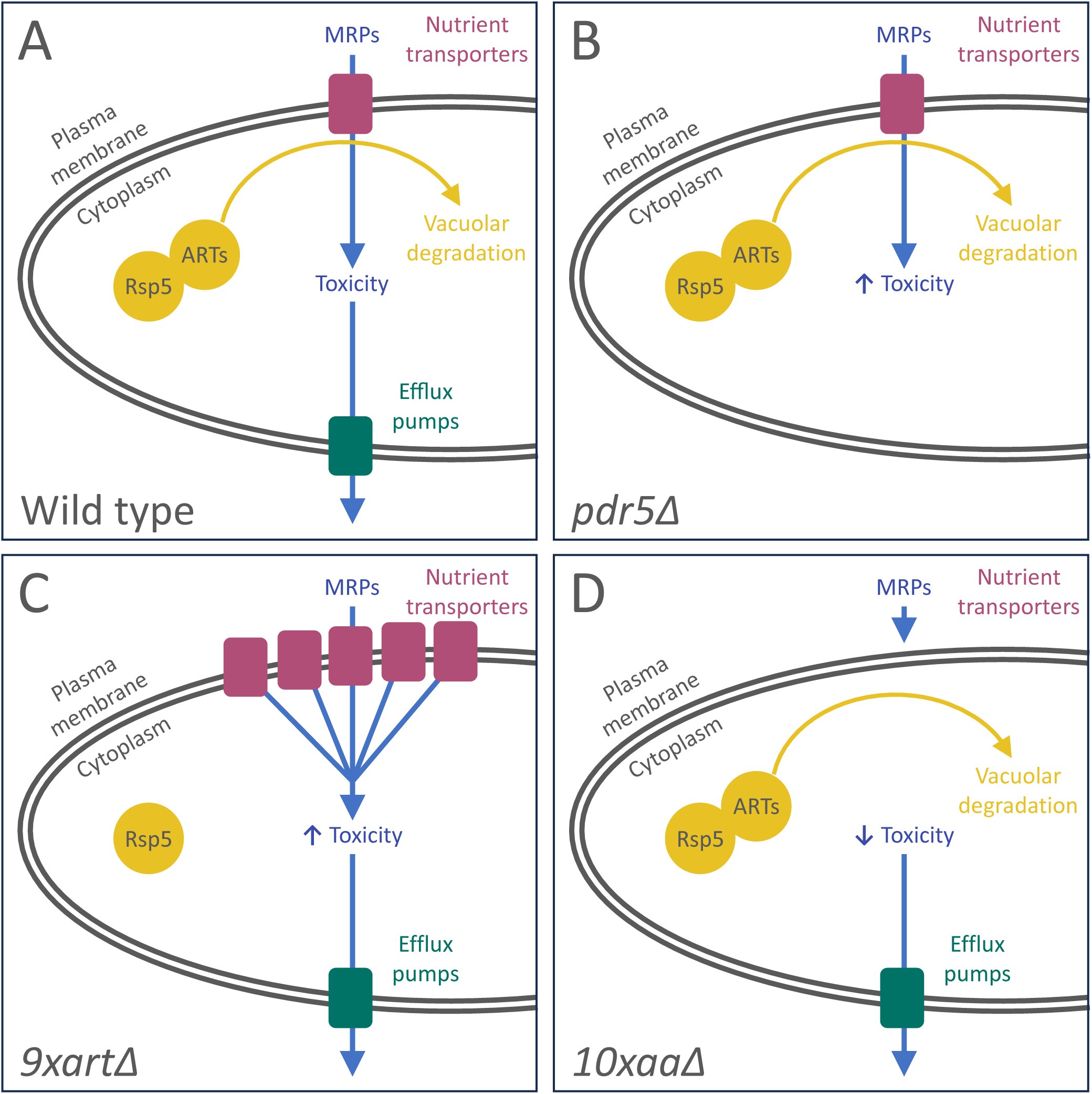
Genetic systems to test sensitivity of furanic compounds in yeast **A)** Wild-type yeast expressing nutrient transporters uptake a range of nutrients across the plasma membrane, with toxic compounds being driven out of the cell through the action of efflux pumps like Pdr5. **B)** If harmful compounds, such as furanic compounds, are substrates of the Pdr5-efflux pump, an increase in toxicity would be predicted in *pdr5Δ* deletion mutants. **C)** Nutrient transporters are routinely downregulated by the vacuolar degradation pathway, where the E3 ubiquitin ligase Rsp5 is recruited to specific transporters via ART adaptors to ubiquitinate and trigger their degradation. A strain lacking nine of these ART adaptors (*9xartΔ*) has higher levels of different transporters at the plasma membrane, which we hypothesize would elevate toxicity via uptake of any compounds by transporters. **D)** Conversely, if toxic compounds rely on transporters for uptake into the cell, a strain lacking many different amino acid transporters (*10xaaΔ*) would be protected from the effects.

### The Pdr5 efflux pump selectively regulates furanic compounds

Pdr5 is a multidrug surface transporter that drives efflux of many toxic molecules to confer resistance (Balzi et al., 1994; Leppert et al., 1990). We reasoned that the harmful effects of high doses of different furanic compounds could be ameliorated by Pdr5, predicting *pdr5Δ* cells would be hypersensitive to MRPs (**Figure 3B**). To test this hypothesis, we compared growth of wild-type yeast and *pdr5Δ* deletion mutants in the presence of previously identified toxic concentrations of HMF, FUR and FMK. Although wild-type and *pdr5Δ* mutants have no significant growth difference in DMSO control media, we found *pdr5Δ* cells are hypersensitive to HMF, growing to only 68% ± 9 of wild-type cells in the presence of 250 μg/mL HMF (**Figure 4A**). At the higher concentration of 500 μg/mL HMF, wild-type cells grow to 88% ± 10 but *pdr5Δ* cells 37% ± 7. Experiments with FUR at concentrations that inhibit growth showed no significant difference between wild-type cells and mutants with *PDR5* deleted (**Figures 4B**). 125 μg/mL FUR reduced growth of wild-type cells and *pdr5Δ* cells to 60% ± 11 and 54% ± 16, respectively. 250 μg/mL FUR did exert a more potent effect in wild-type cells 21% ± 5, but was very similar to *pdr5Δ* cells that retained only 19% ± 10. In much the same way, both 125 and 250 μg/mL FMK inhibited growth of wild-type and *pdr5Δ* cells in a similar, albeit concentration dependent manner (**Figure 4C**). We speculate that wild-type cells might efflux HMF via Pdr5, resulting in the growth benefit of wild-type cells compared to *pdr5Δ* mutants. These results further suggest that each furanic compound affects yeast cells through a different mechanism: HMF appears to be actively effluxed by Pdr5 at toxic concentrations, whereas FUR and FMK show no evidence of Pdr5-dependent transport. Therefore, the cellular response to these compounds is not uniform but compound-specific. Internal control experiments were included in all experiments, showing that wild-type and *pdr5Δ* strains reached a similar final cell density in media treated with DMSO alone.

**Figure 4:**
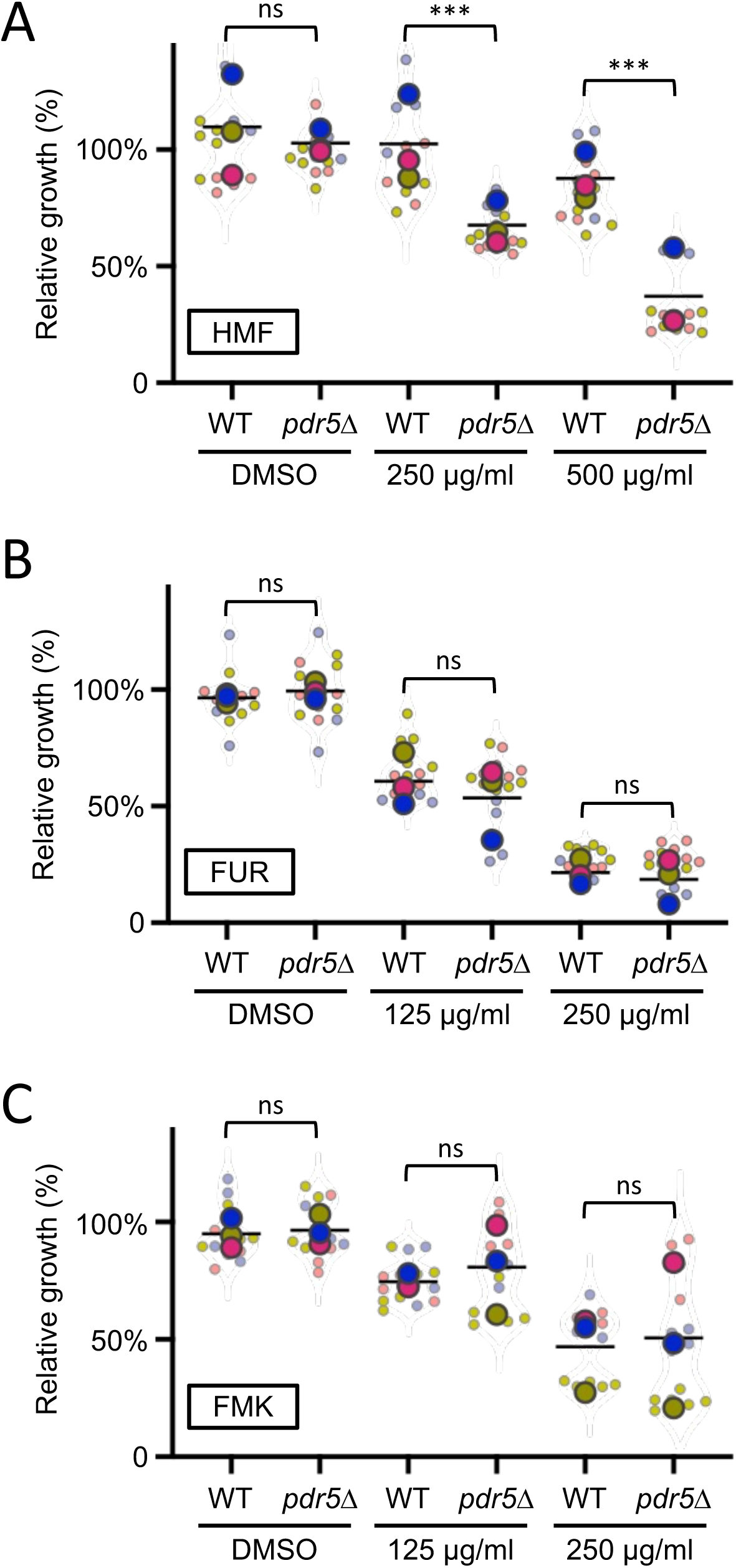
*pdr5Δ* mutants are hypersensitive to exposure of HMF **A, B, C)** Relative growth of wild-type (WT) and *pdr5Δ* cells was assessed following exposure to furanic compounds, with DMSO as a control. Cultures were used to inoculate media containing indicated concentrations of HMF **(A)**, FUR **(B)**, and FMK **(C)** followed by growth measurements after 24 hours at room temperature. One-Way ANOVA and Sidak’s multiple comparisons tests were performed. (ns) not significant, (***) p<0.001.

### Modulation of surface transporters correlates with furanic compound sensitivity

As furanic compounds are relatively similar to natural substrates of yeast transporters, such as amino acids, we considered that nutrient transporters might uptake furanic compounds. To broadly test this idea, we employed a haploid mutant strain lacking 9 different alpha-arrestin (*9xart*Δ) or arrestin-related trafficking (ART) adaptors (Nikko & Pelham, 2009). These proteins act to bridge an array of nutrient transporters with the promiscuous E3 ubiquitin ligase, Rsp5 (Kahlhofer et al., 2021). Deletion of arrestins therefore elevate surface levels of nutrient transporters, allowing increased uptake of exogenous molecules, potentially including furanic compounds (**Figure 3C**). There was almost no difference in growth between wild-type and *9xart*Δ cells upon addition of HMF after 48 hours at either concentration, besides a very subtle hypersensitive phenotype at 500 µg/ml HMF in *9xart*Δ (**Figure 5A**). Much more strikingly, the *9xartΔ* strain was hypersensitive to FUR, with concentration dependent growth inhibition at both 250 and 500 μg/mL concentrations (**Figure 5B**). Similar to HMF, different doses of FMK were equally inhibitory between wild-type and *9xartΔ* strains (**Figure 5C**). These data suggest FUR, and potentially HMF when administered at higher doses, may require surface localisation of nutrient transporters to exert toxic effects on yeast cells.

**Figure 5:**
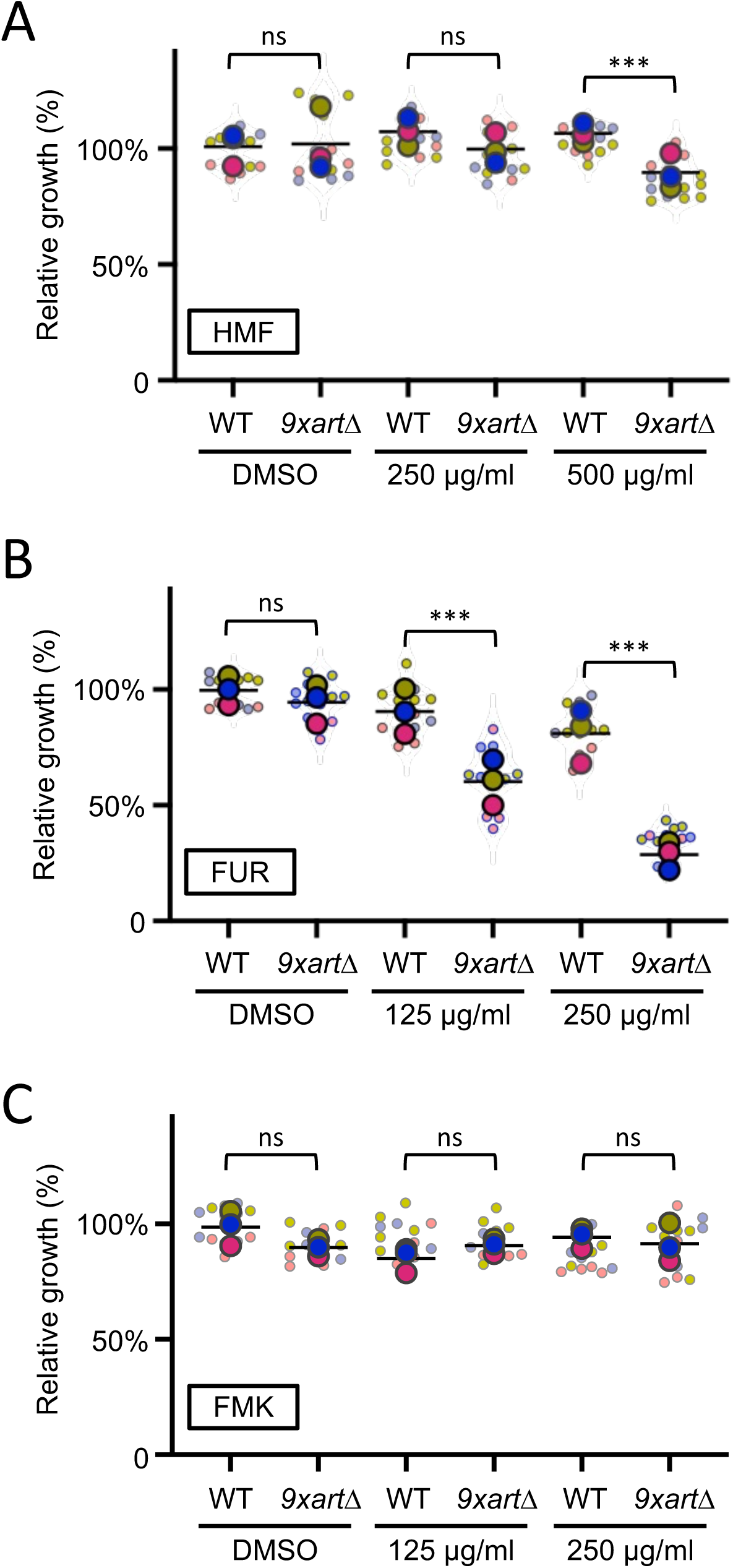
Perturbation of endosomal trafficking adaptors sensitizes cells to furanic compounds **A, B, C)** End point growth assays were used to compare growth in DMSO controls or following exposure to furanic compounds: HMF **(A)**, FUR **(B)**, and FMK **(C)** after 48 hours. Wild-type cells were compared to a *9xartΔ* mutant strain across all conditions. One-Way ANOVA and Sidak’s multiple comparisons tests are indicated: (ns) not significant, (***) p<0.001.

To further investigate the hypothesis that nutrient transporter retention at the plasma membrane drives MRP-induced hypersensitivity in *9xartΔ* cells, we pursued a complementary experimental strategy. Given that arrestin deletion stabilizes transporters at the cell surface, we reasoned that eliminating the transporters themselves would yield the opposite phenotype (**Figure 3D**). To test this, we used the *10xaaΔ* strain; this strain lacks 10 distinct nutrient transporters and is severely deficient in the uptake of most proteinogenic amino acids (Besnard et al., 2016). These experiments revealed that the *10xaaΔ* strain is more resistant to HMF and FUR than wild-type cells (**Figure 6A - 6B**), but reaches a similar final cell density as wild-type cells under DMSO control conditions. This pattern is consistent with the possibility that HMF and FUR exert reduced inhibitory effects when major surface nutrient transporters are absent. This interpretation is further supported by the observation that these compounds elicit hypersensitive phenotypes in mutants with elevated surface transporter abundance (**Figure 5**). Furthermore, FMK, which had no arrestin-related phenotype, also does not have any significant changes in effects in wild-type cells compared with the *10xaaΔ* strain (**Figure 6C**). Taken together, these observations suggest that, within the conditions tested, furanic compounds are actively uptake and processed in a selective manner by nutrient transporters at the yeast plasma membrane.

**Figure 6:**
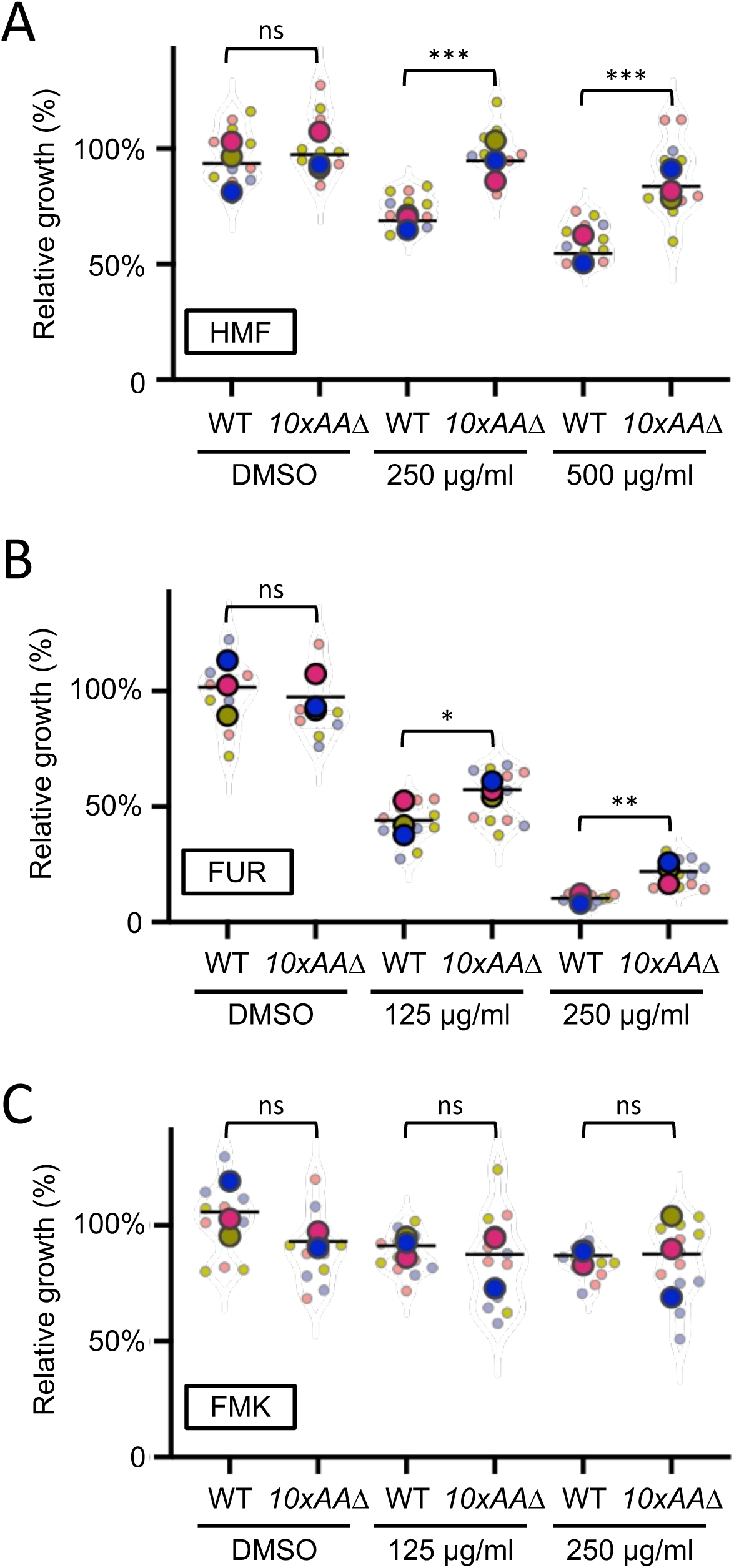
Cells lacking nutrient transporters tolerate furanic compounds **A, B, C)** Relative growth of wild-type (WT) cells was compared to a mutant lacking 10 different amnio acid transporters (*10xaaΔ*). Cells were exposed to HMF **(A)**, FUR **(B)**, and FMK **(C)**, with DMSO added as a control for 24 hours porior to measurements. One-Way ANOVA and Sidak’s multiple comparisons tests were performed. (ns) not significant, (*) p<0.05, (**) p<0.01, (***) p<0.001.

### Furanic compounds perturb distinct cellular processes

We show that yeast can be used to study MRP effects on eukaryotic cells, and that MRPs like HMF, FUR, and FMK appear to exert their effects through distinct mechanisms. To support this idea, we surveyed broad biological processes through organelle maintenance in response to furanic compounds, to indicate if specific pathways were involved in recognising or responding to these MRPs. Firstly, as the Maillard reaction is associated with oxidative stress and mitochondrial dysfunction (Allen et al., 2010; Kim & Hahn, 2013; Qiu et al., 2022), we assessed mitochondrial morphology using a GFP tagged version of Tom6, which localises to a ribbon morphology in wild-type cells (Shaw & Nunnari, 2002). We observed a striking defect in mitochondrial morphology following exposure to HMF for 4 hours, with GFP signal restricted to small bright puncta within the cell (**Figure 7A**). Several mis-localisation patterns of GFP-Tom6 were documented in the presence of HMF, which were absent from DMSO control treatments (**Figure S1A**). Mitochondria span large regions of the yeast cells, so we performed 3D confocal microscopy to better appreciate the morphology in control cells, which show contiguous ribbons (**Supplemental Movie S1**). 3D imaging of the same GFP-Tom6 expressing cells treated with HMF revealed punctate localisations spread throughout the cytoplasm of the cell (**Supplemental Movie S2**). We quantified cells across treatments as a percentage of the total population that exhibit normal, ribbon like mitochondrial structures, with only HMF exhibiting a significant defective morphology phenotype (**Figure 7B**). We then combined HMF treatment with the hypersensitive *pdr5*Δ strain and assessed mitochondrial membrane potential (ΔΨ*_m_*) using MitoTracker Red CMXRos (Poot et al., 1996). These stained mitochondria also showed fragmentation following HMF treatment, suggesting GFP-Tom6 was a faithful representation of mitochondria (**Figure 7C**). The signal intensity of MitoTracker was used as a proxy for ΔΨ*_m_*, to show a significant decrease following HMF treatment, suggesting impaired mitochondrial function (**Figure 7D**).

**Figure 7:**
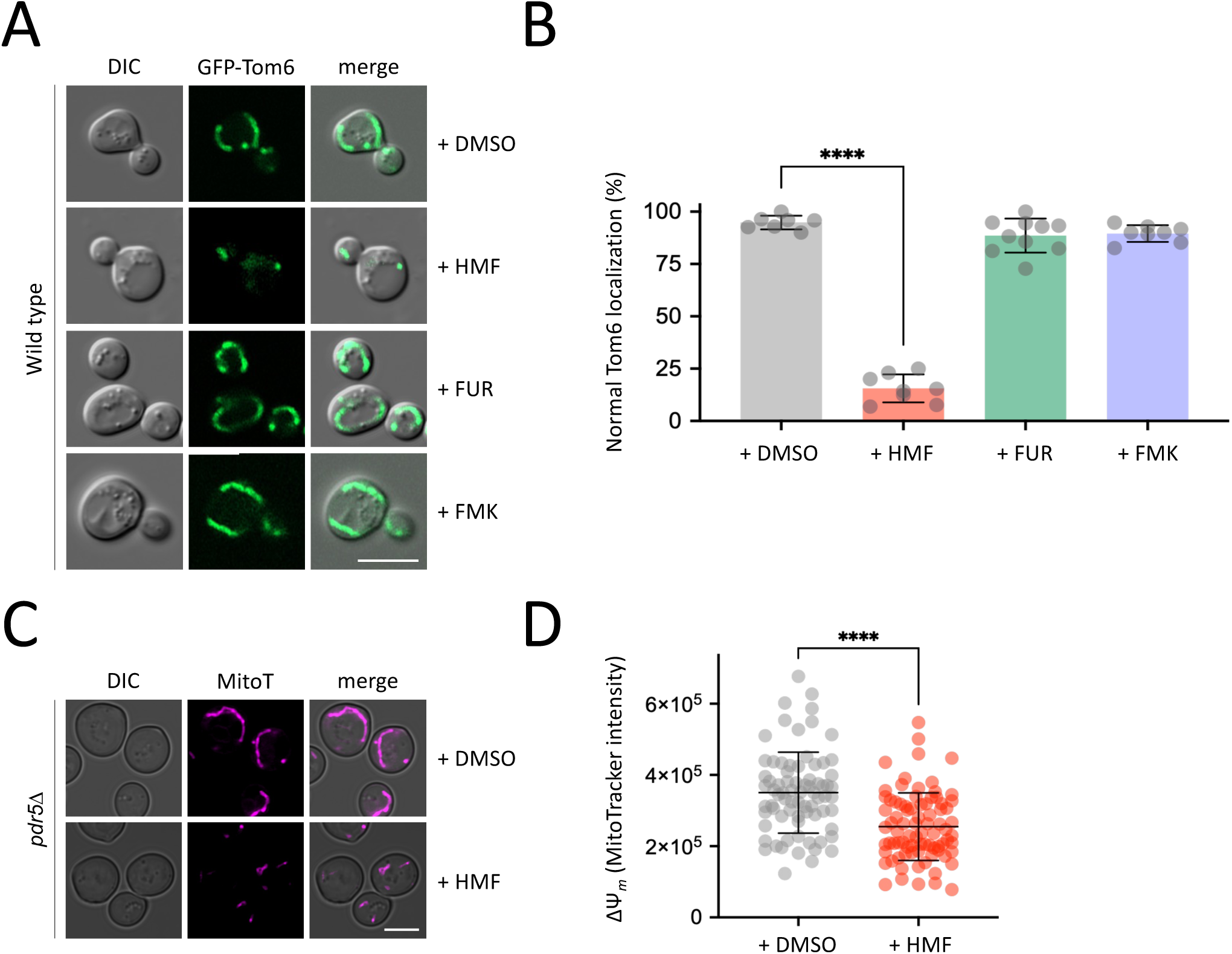
HMF exposure induces mitochondrial defects **A)** Wild-type cells expressing a GFP tagged version of Tom6 (GFP-Tom6) were grown to mid-log phase and imaged by fluorescence microscopy following 4 hours exposure to HMF, FUR, FMK and DMSO as a control. **B)** The average number of cells (n = 150 - 200, per condition) across replicates (n > 6) that exhibit defective mitochondrial morphology are presented. **C)** Mid-log *pdr5Δ* cells were grown to mid-log phase, treated with DMSO or 2000 µg/ml HMF for 4 hours prior to labelling of mitochondria using 2 µM MitoTracker CMXRos. Cells were imaged using laser scanning confocal microscopy. **D)** The fluorescence intensity of MitoTracker stained cells from (C) was measured as an indirect measure of mitochondrial membrane potential (ΔΨ*_m_*), with n > 75 cells measured per condition. (\**\*\*\**) p < 0.0001 shown by unpaired Holm–Sidak *t*-test. Scale bars: 5 µm.

Secondly, as our genetic perturbation of nutrient transporters and arrestins correlated with sensitivity to furanic compounds, we considered the endolysosomal system as a potential target organelle. We chose the vacuole as a representative marker, as it is also known to respond to different stress conditions (Kohler & Büttner, 2021; Li & Kane, 2009). Cells expressing a GFP tagged version of the vacuolar ATPase subunit Vph1 were grown to mid-log phase and then exposed to inhibitory concentrations of different furanic compounds for 4 hours. As expected, the vacuoles of wild-type cells exposed to DMSO showed no defects, which was also true of HMF or FMK treatments (**Figure 8A**). However, FUR induced a variety of abnormal phenotypes never observed in control cells, including significant GFP-Vph1 inside the lumen of the vacuole, in addition to accumulations at the periphery, alongside other non-vacuolar signal (**Figure S1B**). Cells were quantified as a percentage exhibiting normal GFP-Vph1 localisations, with FUR treatment the only condition that caused a significant number of cells with abnormal localisations (**Figure 8B**). This vacuolar disruption phenotype was corroborated in hypersensitive *9xart*Δ cells using the lipid dye FM4-64 following a pulse-chase protocol to label the vacuole, with cells treated with either DMSO or FUR (**Figure 8C, S1C**). Again, FUR induced a significant percentage of cells with an obvious intralumenal signal of FM4-64 compared to DMSO treatment, where the signal is exclusively localised to the limiting membrane of the vacuole (**Figure 8D**).

**Figure 8:**
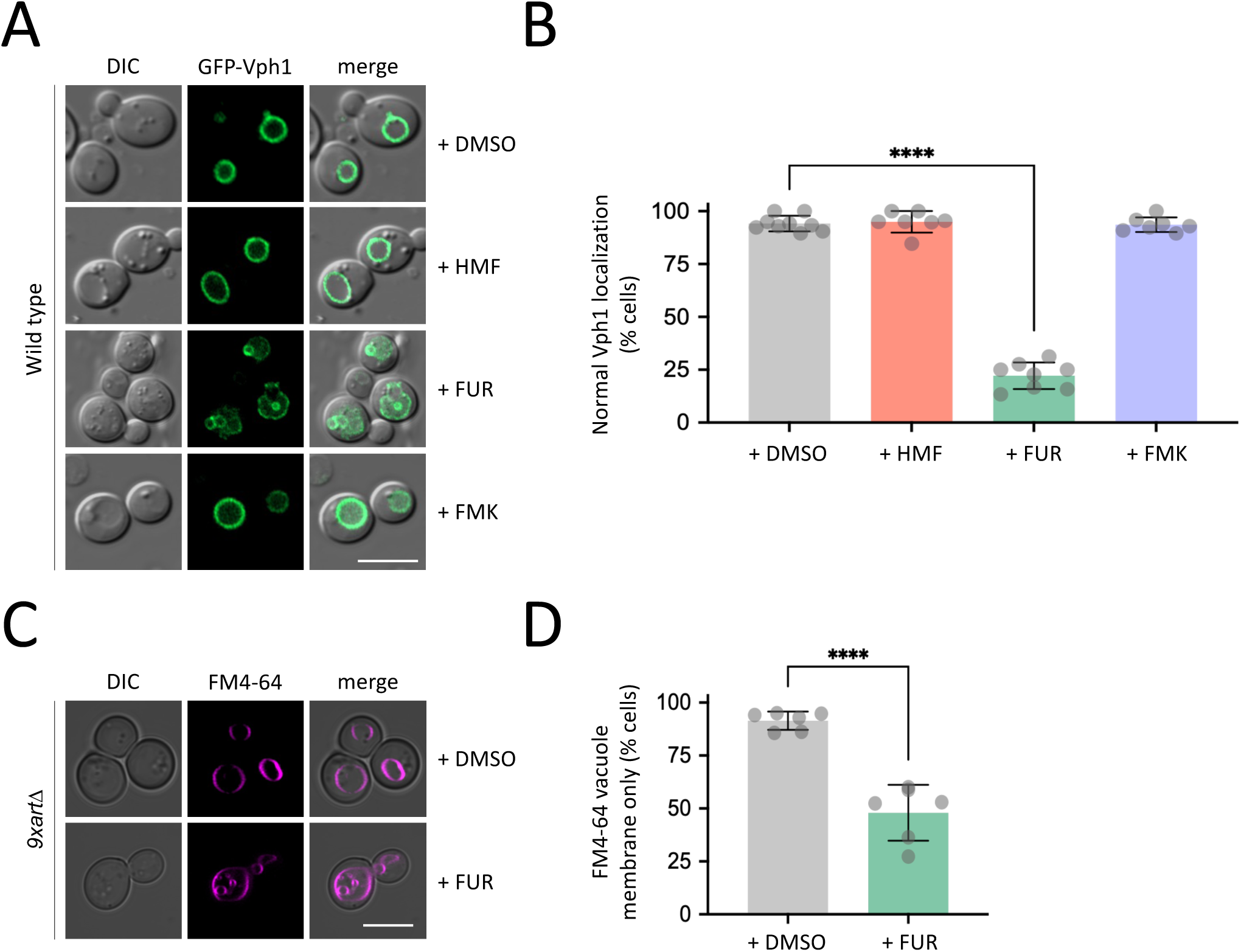
Vacuolar morphology is perturbed following exposure to FUR **A)** Cells expressing GFP-Vph1 were grown to mid-log phase prior to DMSO or furanic compounds (HMF, FUR and FMK) being added to the media for 4 hours prior to confocal microscopy. **B)** The average number of cells (n > 150 per condition) across replicates (n > 5) that exhibit abnormal vacuolar localisations are presented in the histogram as a percentage. **C)** *9xart*Δ cells were grown to log phase prior to 4-hour exposure 1000 µg/ml FUR, with control cells treated with DMSO for the same period. Cells were then labelled with 0.8 µM FM4-64 for 30 minutes, followed by a 1-hour chase period in label free media. Pulse chase steps in the protocol were performed with DMSO and FUR supplemented media. Labelled vacuoles were then imaged by fluorescence microscopy. **D)** Cells from (C) were quantified from experiments (n > 5) and the percentage of cells exhibiting only FM4-64 labelling at the limiting membrane was calculated, compared to cells that were observed to also have intravacuolar signal. (\**\*\*\**) p < 0.0001 as determined by unpaired Holm–Sidak *t*-test comparing treatments to DMSO control. Scale bars: 5 µm.

This physiological analysis further supports the idea that individual MRPs have cell specific responses and sensitivities in eukaryotic cells and that many layers of regulation are likely involved in any cellular response to even individual compounds.

## DISCUSSION

The intermediate Maillard reaction products classed as furanic compounds are chemical contaminants created during industrial processes or cooking, commonly identified in food following excessive heating (e.g. honey and caramelised syrups) or on the surface of fried or baked foods (Alsafra et al., 2022; Conceição et al., 2024; Santos et al., 2025; Shapla et al., 2018; Sun et al., 2022). These compounds can form through sugar degradation, lipid peroxidation, or thermal decomposition of amino acids, indicating their diverse origins and potential variability in concentration depending on food matrix and processing method (Hosry et al., 2025). EFSA CONTAM Panel identified a worrying range of furan exposure, particularly consumption of ready-to-eat meals and cereals for infants and coffee for adults (EFSA Panel et al., 2017). Given their presence in foods of habitual consumption and their constant exposure in the human diet, it is crucial to evaluate the dietary exposure to these compounds and to understand the mechanisms associated with their uptake, efflux, and impact on processes at the cellular level. Several studies have also highlighted the potential for furanic compounds to induce oxidative stress and mitochondrial dysfunction, which can contribute to cytotoxicity and metabolic disruption in exposed cells (Allen et al., 2010; Aydin et al., 2023; Batool et al., 2021, 2025; Zhang et al., 2015). In this study we focussed on three furanic products from the Maillard reaction commonly found in food: HMF, FUR, and FMK, which present different functional groups, such as hydroxyl, methyl, and ketone, in the molecular rearrangement linked to the structure of a furan ring (**Figure 1**). Although these compounds share molecular features, it is important to consider their differences, which might influence cytotoxic potential, mechanisms and specificity of transport in/out of cells, and mode of action that perturbs cellular processes. Moreover, differential interactions with membrane transporters and detoxification enzymes may underlie the distinct cellular responses observed for each compound, emphasising the need for compound-specific toxicokinetic studies (Quan et al., 2022) . Understanding these nuances is key to assessing the risk associated with dietary intake of Maillard reaction products and developing strategies to mitigate their impact on human health.

Although previous studies have reported toxic effects of furanic compounds on the growth of eukaryotic cells, including yeast (Ask et al., 2013; Lam et al., 2021), our documentation of a dose response inhibition on growth in response to HMF, FUR and FMK suggests regulatory mechanisms are triggered by this stress, and provide optimised concentrations to explore these physiological responses. As mentioned, although MRPs are complex and often conjugated to other molecules, studying MRPs in isolation best represents the common products encountered by cells (Pagare et al., 2024). Curiously only HMF showed clear phenotypes suggesting clearance via the Pdr5 multidrug resistance pump, with *pdr5Δ* mutants being hypersensitive to different concentrations (**Figure 4**). This specificity may be related to the higher polarity or redox activity of HMF compared to FUR and FMK, which could enhance recognition or affinity to the substrate-binding cavity for efficient Pdr5 efflux (Egner et al., 1998). As a fungal protein, much of the work on Pdr5 is focussed on biomedical and agricultural applications (Golin & Schmitt, 2023). However, this family of ATP-binding cassette (ABC) proteins are conserved from bacteria to humans (Balzi et al., 1994), with various human homologues shown to transport a range of structurally and functionally distinct substrates across the plasma membrane (Lin & Yamazaki, 2002; Sodani et al., 2012). Therefore, we suggest that multidrug resistance pumps are worthy of consideration for regulation of MRPs consumed in human diets.

We do note that the genetic systems we have used to understand the tolerance of yeast to furanic compounds rely on indirect assumptions. It is possible that, although we know more/fewer nutrient transporters localise and function at the plasma membrane in *9xartΔ* and *10xaaΔ* cells, respectively (Besnard et al., 2016; C. H. Lin et al., 2008; Nikko et al., 2008; Nikko & Pelham, 2009). Alternative explanations could be through the intracellular targets of furanic compounds being mis-localised in these mutant cells, or other indirect consequences, such as membrane destabilisation under stress (Teixeira et al., 2008) . However, given results such as *9xartΔ* cells are hypersensitive to FUR and *10xaaΔ* mutants are resistant, we believe the most logical explanation is transport via a surface localised transporter facilitating cellular entry of FUR. To extend this further, we assume at least one of the 10 transporters deleted in the *10xaaΔ* strain has elevated surface retention in the *9xartΔ* strain. Although we acknowledge the nature of what transporter(s) are responsible for these responses is not yet known, and yeast transporters are known to vary in their substrate specificity (Bianchi et al., 2019). Furthermore, environmental pressure can modify yeast transporters, either modulating specificity or creating new activity (Karapanagioti et al., 2024). Therefore, it is reasonable to assume that these furanic compounds, which are chemically similar to natural yeast transporter substrates, might utilise transporter activity to enter cells. Changes in exogenous substrate levels, such as glucose or nitrogen depletion, indirectly regulates surface transporters via post-translational modification of arrestins (Kahlhofer et al., 2021; Megarioti et al., 2021; O’Donnell & Schmidt, 2019). Therefore, as proteins like amino acid transporters and alpha arrestins are highly conserved with an array of human orthologues that perform analogous functions in animal cells (Gauthier-Coles et al., 2021; Zbieralski & Wawrzycka, 2022), these uptake mechanisms of furanic compounds may be conserved in other eukaryotic systems.

Our transporter-based phenotypes provide a mechanistic counterpoint to the longstanding assumption that furanic aldehydes enter cells exclusively by passive diffusion. This view, rooted in classical fermentation literature, has been widely accepted due to the small size and modest polarity of HMF and furfural. However, the hypersensitivity of *9xartΔ* mutants, where nutrient transporters accumulate at the plasma membrane, and the striking resistance of *10xaaΔ* strains lacking major amino acid permeases reveal that transporter-mediated uptake contributes substantially to intracellular accumulation of FUR and, at higher doses, HMF. Thus, our data support a dual-entry model in which passive diffusion provides a basal route, while surface transporters amplify toxic exposure when dysregulated. This framework reconciles historical diffusion-based models with modern transporter biology and suggests that similar transporter-dependent effects may be conserved across eukaryotes. Our results can also be used to contextualise previous efforts to understand toxic effects of furanic compounds, a common issue during conversion of lignocellulosic material for biofuel and chemical production (Jönsson et al., 2013). A CRISPR screen has previously been performed to identify yeast genes involved in tolerating lignocellulosic toxins, including the Ume6 transcriptional regulator and the Hog1 stress response gene (Gutmann et al., 2021). Ume6 has been shown to indirectly regulate nutrient transporter trafficking (Amoiradaki et al., 2021) and Hog1 activity is thought to correlate with expression of surface transporters in response to osmotic stress (Guo et al., 2026), further suggesting that toxicity could be regulated by active transport.

The intracellular targets of internalised furanic compounds are not well understood, but physiological effects reported in animal cell models have been proposed in the context of mitochondrial dysfunction (Qiu et al., 2022). To test this in yeast, we expressed a GFP-tagged version of Tom6, a component of the Translocase of the Outer Mitochondrial membrane complex (Pfanner et al., 1996). Under control conditions with DMSO, or even with addition of FUR and FMK at concentrations that inhibit growth, we observed no obvious defects in mitochondria morphology, even using 3D confocal microscopy. In contrast, addition of HMF induced fragmentation of mitochondria, with foci dispersed throughout the 3D volume of the cytoplasm (**Figure7**). This suggests furanic compounds drive increased mitochondrial fission, an adaptive stress response in eukaryotic cells associated with starvation and stress conditions that can lead to cell death (Pagliuso et al., 2018; Wang et al., 2022; Westermann, 2012). We also observed a decrease in mitochondrial activity following the addition of HMF. Thus, changes in mitochondrial morphology and activity in yeast may correlate with conserved oxidative stress and metabolic and inflammatory responses following furanic compound exposure in animal cells (Lee et al., 2019; Qiu et al., 2022; Shapla et al., 2018). These observations also align with research showing exposure of furanic compounds to different parasite species, including *Trypanosoma cruzi* and *Leishmania amazonensis*, inhibits growth (Hartmann et al., 2017; Sifontes-Rodríguez et al., 2015). Indeed, a recent study in the parasite *Toxoplasma gondii* showed growth defects and mitochondrial perturbations following exposure to furanic compounds (Portes et al., 2025). Curiously, no obvious mitochondrial defects of FUR and FMK were observed, at the concentrations tested. To further explore potential targets of these other furanic compounds, we considered the vacuole a potential target given the data implicating transporters and endolysosomal trafficking (**Figure 5 &** 6). Furthermore, the vacuole is associated with downregulation of transporters in response to other stresses, like different nutrient starvation conditions (Laidlaw et al., 2021; Müller et al., 2015; Paine et al., 2021). Although HMF had no effect on vacuolar morphology, FUR induced a series of mis-localisation phenotypes, that may be directly or indirectly related to the stress induced by FUR leading to cell death (**Figure 8**).

Although FMK concentrations that effectively inhibit yeast growth in a concentration manner were established, our genetic models did not identify conditions where cellular sensitivity or resistance to FMK were altered compared to wild-type controls, and FMK did not exhibit morphological defects in mitochondria or vacuoles. This suggests that FMK may provide toxicity by an entirely distinct mechanism, or a redundant combination of mechanisms. Less is known about FMK in the literature but this compound has already been identified in several thermally processed foods and can inhibit microbial driven production of biofuel (EFSA Panel et al., 2017; Guarnieri et al., 2017; Sayre et al., 1993). Beyond this, structurally related methylfuran compounds have been shown to form adducts with essential cytosolic proteins and perturb NAD⁺/NADH balance in yeast (Jilani & Olson, 2023), indicating FMK might similarly disrupt central metabolism rather than organellar integrity.

Having identified concentrations to study FMK in yeast, future work may provide insight as to its mode of action, which would be relevant for understanding toxic effects in humans. In conclusion, MRPs are complex and diverse, so concentrating our study on not just one but three furanic compounds allowed the specificity and modes of action to be compared. Strikingly, even across relatively similar compounds HMF, FUR and FMK, when studied in isolation we find each elicits differences: in effective concentrations, specificity for uptake/efflux, and perturbation of distinct organelles. The extensive complexity of these physiological responses will be difficult to unravel, but our work provides a framework to do so in model organisms that can reveal novel and relevant mechanistic models that can be tested in animal cell models. Beyond this, as such MRPs are inhibitory to biotechnological processes, such as the production of biofuel (Allen et al., 2010; Ask et al., 2013; Ren et al., 2024; Wang et al., 2016), this study can guide applications to modulate yeast tolerance for improved processing. Our findings contribute to a better understanding of how even relatively similar furanic compounds elicit different organelle-specific effects. This complexity that should be considered when trying to define the effects of MRPs in the context of human health. Elucidating these mechanisms in yeast not only facilitates screening for dietary safety thresholds but also aids in identifying biomarkers of exposure or damage that could be relevant for human toxicology. Mechanistic knowledge strengthens food risk assessment frameworks and supports evidence-based regulation of processing contaminants.

## MATERIALS & METHODS

### Yeast strains used in this study

Parental yeast strain BY4742 (Brachmann et al., 1998) was used for experimental work throughout the manuscript. BY4742 was the parental strain to test the role of Pdr5 (BY4742 *pdr5Δ::Kan^r^* (Giaever et al., 2002) and alpha arrestins (*9xartΔ*: BY4742 *ecm21Δ csr2Δ bsd2Δ rog3Δ rod1Δ ygr068cΔ aly2Δ aly1Δ ldb19Δ ylr392cΔ* (Nikko & Pelham, 2009). BY4741 (Brachmann et al., 1998) was the parent for the *NOP1*-GFP-Tom6 and *NOP1*-GFP-Vph1 strains for confocal microscopy (Weill et al., 2018; Yofe et al., 2016). To assess the role of yeast lacking nutrient transporters, the Σ22574d parental strain (Jauniaux & Grenson, 1990) was used following deletion of ten distinct transporters (*10xaaΔ*: *gap1Δ put4Δ uga4Δ can1Δ lyp1Δ apl1Δ hip1Δ dip5Δ gnp1Δ agp1Δ* (Besnard et al., 2016).

### Cell culture

Prior to culturing, yeast strains were grown on yeast extract peptone dextrose (YPD) agar medium plates (2% peptone, 2% glucose, 1% yeast extract, 2% agar) (Formedium, Norfolk, UK) overnight at 30°C. Yeast cultures were grown in synthetic complete (SC) medium (2% glucose, yeast nitrogen base supplemented with a complete mixture of required amino acid, bases and vitamins) (Formedium, Norfolk, UK) overnight at 30°C with shaking to early/mid-log phase (OD_600_ ≤ 1.0) prior to experimentation.

### Steady state growth assays

WT yeast cultures were grown overnight to saturation and 1.5 µl of culture was used to inoculate 200 µl SC medium with a stated titration of three furfural compounds (FC): 5-Hydroxymethylfurfural (HMF)), Furyl methyl ketone (FMK) or Furfual (FUR) (Sigma-Aldrich, USA) or Dimethyl sulfoxide (DMSO) was used as a control in a 96-well plate format. Plates were incubated at room temperature and subjected to readings at OD_600_ at 1hr and 24hrs (*pdr5Δ and 10xaaΔ*), and 1hr and 48hrs (*9xartΔ*) using a microplate spectrophotometer (ThermoMultiskan Go 1510 Sky, Thermo Fisher Scientific Inc., Massachusetts, US). Data was normalized to % cell viability, considering the average growth of each cell strain (without any exposure) as 100% and plotted in GraphPad Prism (version 8) to compare the statistical significance between experimental conditions, with *P*-values included. All end point assays are presented from at least 6 technical and 3 biological repeats. An asterisk (*) is used to denote significance *p* < 0.05.

### Continuous growth assays

96-well plates were set up following the same method as above. Growth experiments were monitored continuously for 24 hrs using a Stratus microplate reader (Cerillo, Charlottesville, VA, US) at 30°C. Growth curves presented are averages of at least six technical replicates.

### Confocal microscopy

GFP-labelled yeast strains were grown overnight to mid-log phase, cells were pelleted and resuspended in media supplemented with the three different FC compounds at stated concentrations. For experiments using fluorescent dyes, mid-log phase cells were first cultured prior to labelling. For mitochondria straining, 2 µM MitoTracker CMXRos (Thermo Fisher Scientific) was incubated for 30 minutes at 30°C, then washed. For vacuolar staining, 0.8 µM (*N*-(3-Triethylammoniumpropyl)-4-(6-(4-(Diethylamino) Phenyl) Hexatrienyl) Pyridinium Dibromide) FM4-64 (Thermo Fisher Scientific) and incubated for 30 minutes, followed by a 1-hour chase period in label free media. All media used for labelling, washing and chasing, for either MitoTracker or FM4-64, was supplemented with DMSO or stated furanic compound. Cells were imaged at room temperature using a 63x 1.40 oil immersion Plan Apochromat objective lens on a laser scanning confocal microscope (Zeiss LSM980). GFP fluorescence was excited using a 488 nm Argon laser and 495–550 nm emission collected. For cells labelled with red fluorescent dyes, imaging was performed at room temperature using a 63x 1.40 oil immersion Plan Apochromat objective lens on a Zeiss LSM 980 with Airyscan2. MitoTracker red and FM4-64 fluorescence was excited using a 561 nm Argon laser and 570-620 nm emissions collected. Processed images were adjusted in Fiji/ImageJ software (NIH).

## Statistical analyses

One-Way ANOVA followed by Sidak’s multiple comparisons test was utilized to compare wild-type (WT) cells with deletion strains under the same conditions to determine significant differences (p < 0.05). Additionally, an unpaired *t-*test was performed to compare normal organelles from GFP-exposed cells with control cells (GraphPad Prism v8, USA).

## ACKNOWLEDGMENTS

We would like to thank staff at the York Bioscience Technology Facility for technical assistance. We are very grateful to Guillaume Pilot (Virginia Tech) and Foteini Karapanagioti for providing the *10xaaΔ* yeast strain for experimental work. This research was supported by a Sir Henry Dale Research Fellowship from the Wellcome Trust and the Royal Society 204636/Z/16/Z (CM) and by the National Council for Scientific and Technological Development (CNPq-Brazil) 405816/2021-9 (LCPM).

