## Supplementary figures and images for "Uptake mechanisms and physiological effects of furanic compounds from the Maillard reaction in budding yeast"

### Supplemental figures

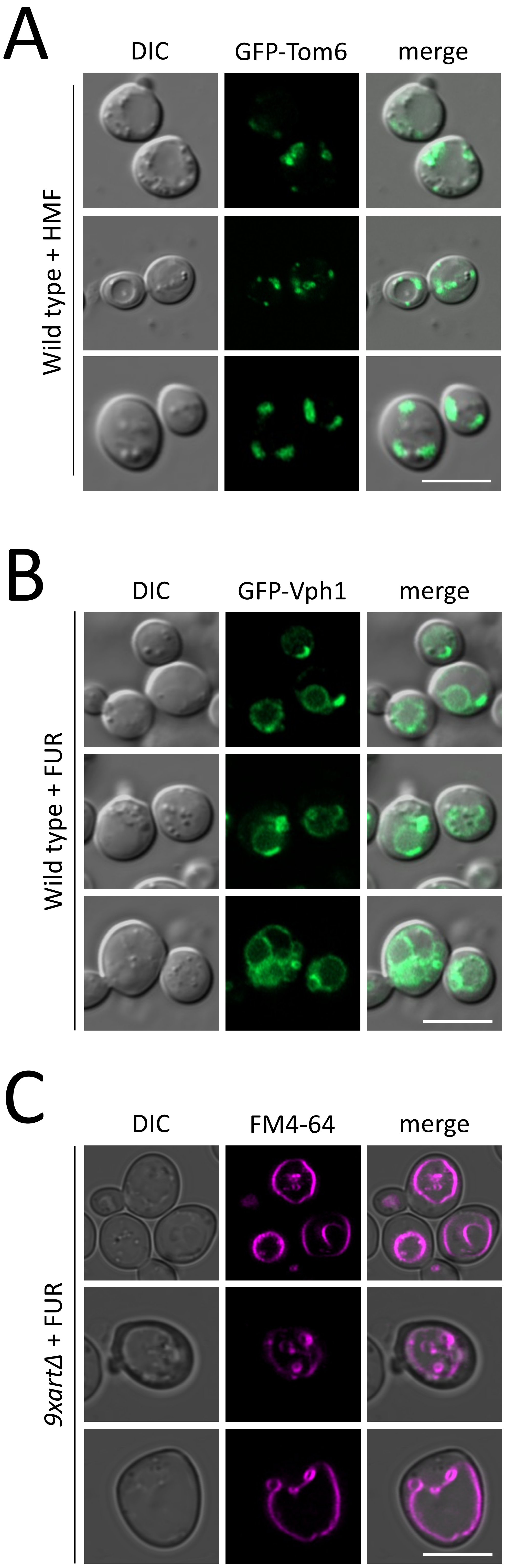
